# Visual awareness during the attentional blink is determined by representational similarity

**DOI:** 10.1101/2022.10.25.513789

**Authors:** Matthew F. Tang, Kimron L. Shapiro, James T. Enns, Troy A.W. Visser, Jason B. Mattingley, Ehsan Arabzadeh

**Author notes:** **Corresponding author** Matthew F. Tang, Eccles Institute of Neuroscience, John Curtin School of Medical Research, The Australian National University, Acton, ACT, 2601, Australia.

## Abstract

Our visual perception seems effortless, but the brain has a limited processing capacity which curtails the amount of sensory information that can be brought into conscious awareness at any moment in time. A widely studied exemplar of this limitation is the ‘attentional blink’ (AB), in which observers are unable to report the second of two rapidly sequential targets if it appears within 200-500 ms of the first. Despite the apparent ubiquity of the AB effect, its computational and neurophysiological underpinnings have remained elusive. Here we propose a simple computational model of temporal attention that unifies the AB with spatial and feature-based attention. We took a novel, integrative approach involving human psychophysics and functional brain imaging, along with neuronal recordings in mice to test this model. Specifically, we demonstrate that the AB only arises when visual targets have dissimilar representations in the brain but is absent when both targets have the same representation. Similarity in this context can be determined either by elementary features such as edge orientation, or by acquired, high-level factors such as numerical or alphabetical order. In this parsimonious model of the AB, attention to an initial target establishes a perceptual filter that is tuned to its unique representation in the brain. Subsequent items that match the filter remain available for conscious report, whereas those that do not match elude awareness altogether.

## Introduction

One of the most persistently difficult challenges in sensory neuroscience has been to determine the limits on conscious awareness. It is generally understood that humans have limited capacity to bring all information that reaches their sensory receptors to the level of awareness. Attention allows us to prioritize just one or two items toward which we can guide our motor actions, at the expense of all others^1,2^. A prominent example of this limitation occurs when people are asked to monitor a stream of sequential items and report the identity of two target items embedded within the stream. People can typically report the identity of the first item (T1) but not the second (T2) when these items are separated by 200-500 ms, a phenomenon known as the attentional blink (AB)^3–5^. There are currently multiple accounts for the AB, with most arguing that the deficit results from the target presentation rate exceeding the brain’s processing capacity. Here we propose an alternative account, which we call ‘representational enhancement’, which is derived from the same mechanism as spatial and feature-based attention. On this account, the second item within a sequential stream is not missed because of the limited capacity of working memory; rather, it is missed because the act of selecting the first item alters the filter controlling conscious awareness of the second item. That filter is tuned to items that possess a similar neural representation to the first target.

The neural underpinnings of many aspects of selective attention are now relatively well understood. Spatial attention increases neural responses to attended items with a corresponding decrease in responses to items in unattended locations^1,6^. Similarly, if attention is directed to specific features, neurons selective for those features show an enhanced response regardless of the item’s location^7,8^. Attending to one item has predictable effects, such as enhancing perceptual sensitivity for stimuli presented at the attended location and a degradation of sensitivity for items presented elsewhere^9,10^. Collectively, these findings suggest that the visual system has a homeostatic-like allocation of processing resources, such that the neural response is amplified for attended locations or features at the expense of unattended locations and features.

Theories that account for the second-item deficit in an AB stream are generally considered separately from those proposed to account for the effects of spatial- and feature-based attention. One class broadly argues that missing T2 is due to extended processing of T1, resulting in insufficient resources being available for T2^3,4,11^. Another class instead argues the deficit reflects the cost of switching between target and distractor processing^12–14^. A third class posits the AB is caused by over-investment of attention to T1 rather than a limit in processing resources^15,16^. This third class is motivated by findings that show a reduction in the AB when participants’ attention to the task is reduced due to mind wandering^17^ or surrounding distractor items^15^. Unlike theories of spatial attention, neural evidence for specific theories of the AB is relatively scarce. Human brain imaging studies have revealed a decreased late-stage component (P300) associated with missing T2^18^ and a decrease in activity in the retinotopic location of T1^19^ during the AB. As another example, the feature-selective representation of T2 is reduced in the presence of AB^20,21^. While these results provide some information about the neural underpinnings of the AB, no work has directly related the findings to the neurophysiological literature on spatial and feature-based attention. Overall, the three classes of theories seem to explain subsets of the AB features and the neural correlates of the AB in a manner that is distinct from the spatial and feature-based attention literature.

The current work asks whether we can place the AB within the well-established framework of spatial and feature-based attention to create a unified theory. Recall that according to spatial and feature-based theories, attending to one item enhances its neural representation, with a corresponding decrease in neural representations of unattended items. A consequence of this enhancement is that items that are represented similarly will receive enhanced processing whereas those that have dissimilar representations will be suppressed. To explain the AB by the same mechanism, we propose that as T1 is attended for identification, its representation becomes enhanced, with the representational similarity of T1 and T2 determining the strength of the AB. If T2 has a dissimilar representation to T1, then its activity will be suppressed, whereas T2’s representation will be enhanced and accurately reported if it has a similar representation to T1. Note that most AB studies have used English alphanumeric characters as targets, which on average tend to have dissimilar representations from one another. In such studies, therefore, the AB is evident in most trials.

Here we used a wide-ranging, multimodal approach to determine whether attending to T1 causes the brain to establish a filter for similarly represented targets. We combined human psychophysics and brain imaging, mouse imaging and computational modelling, to provide support for this unifying account of the AB that places it within the existing framework of spatial and feature-based attention (the ‘representational similarity’ account).

## Results

### Computational model of attentional blink

We began by creating a formalised model of attention based on representational similarity that is consistent with well-known spatial and feature-based accounts of attention^1,2,7^. We start with a simple model that uses orientation as a representational feature, which allows us to quantify differences in representation between the targets (T1 and T2) in a well-established neural feature space. The model consists of six ‘neurons’, each tuned to a different orientation (0° to 150° in 30° steps). At the beginning of each trial, the population is equally sensitive to all orientations (Figure 1A), but this changes after T1 presentation (Figure 1B). The sensitivity of the channels is adapted by the inverse of their response to the orientation of T1, thus suppressing the non-activated channels. This has the same effect as priming or enhancing the activated channels while not affecting the non-activated channels, and is consistent with how spatial and feature-based attention affects neuronal population activity^7,8,22,23^. T1 therefore creates a filter which allows T2 to generate a large neural response only if its orientation is similar. When the orientation of T2 is different the response will be suppressed. There are many well-known studies and theories suggesting that we only consciously perceive targets if the corresponding neural response exceeds a certain threshold^24,25^. If this threshold is not reached then the global, brain wide ‘ignition’ of activity which corresponds with conscious awareness does not occur.

**Figure 1.**
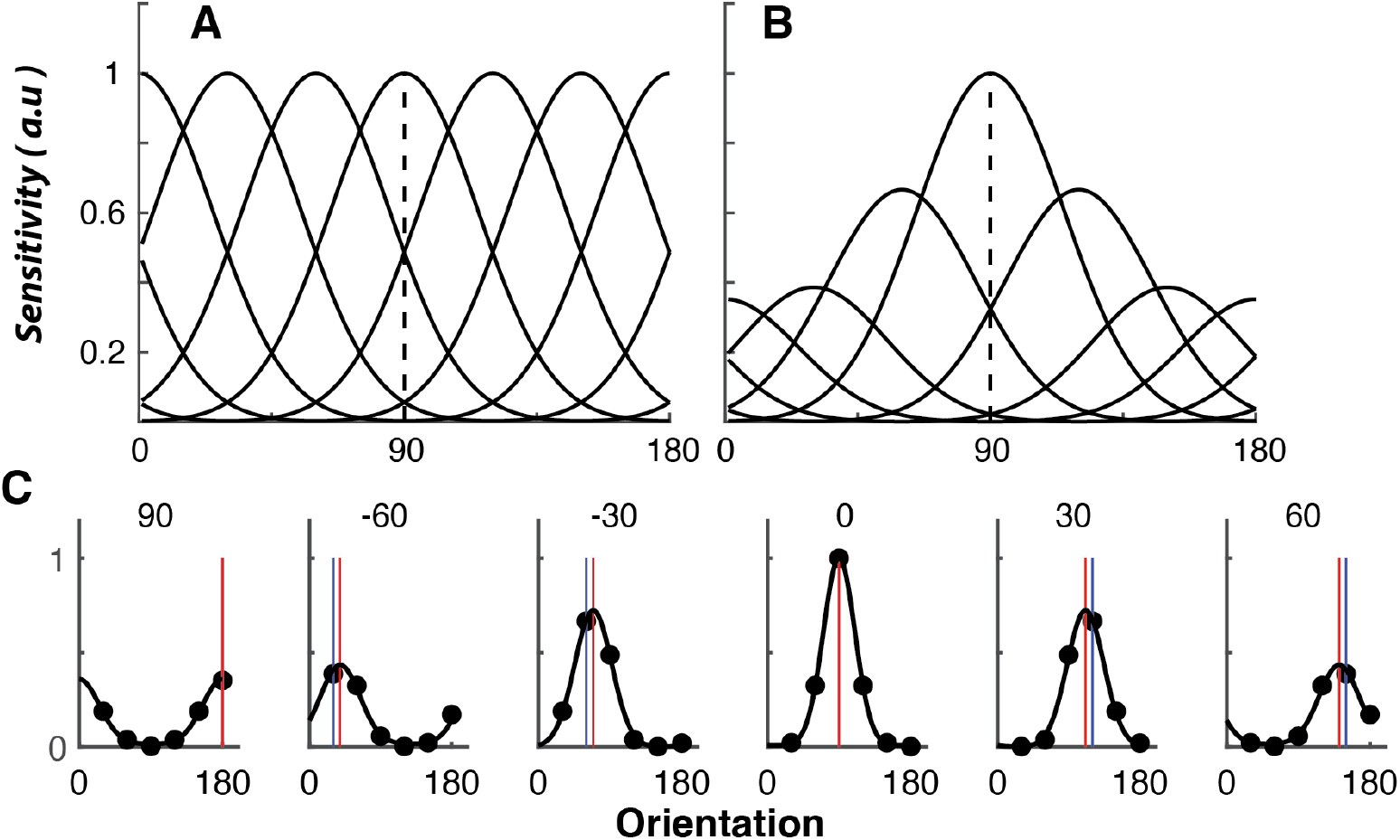
Computational model of stimulus-selective attention to predict perceptual and neural data. The population of neurons is equally sensitive to all orientations before T1 (A) but after a 90° T1 is presented, the selectivity of the channels is enhanced relative to their response (B). (C) The response of the population to T2 depends on the difference in orientation between T1 and T2. The blue vertical line shows the presented T2 orientation and the red line shows the population decoding orientation given by the vector mean. It is evident that the decoded orientation will be biased toward the T1 orientation with the largest biases at 30° to 60° orientation difference between T1 and T2.

In our model, the response of the population to T2 depends on the difference in orientation between T1 and T2 (Figure 1C), with the largest response occurring when the orientations are most similar, and which is gradually diminished with greater orientation differences. This is consistent with psychophysical and neural data showing attention to an orientation has a greater effect on nearby, relative to distant, orientations^26,27^. The resulting T2 response (given by the vector sum of the population response) is also biased toward the T1 orientation. This arrangement predicts that the perceived orientation of the second target will be attracted toward the orientation of the first target. This is consistent with the well-known serial dependency effect, in which the orientation of a target is attracted toward the orientation of the previously-presented target^28,29^.

### Experiments 1 – Conscious awareness of the second target relies on target similarity

We designed a simple visual task to test our representational-similarity model of the AB^20^. Participants (N = 22) were presented with a rapid (8.33 Hz) stream of 20 pseudo-randomly oriented Gabor items and were asked to determine the orientation of the two targets (T1 and T2) which had higher spatial frequencies than the intervening distractor items (Figure 1AB). Using Gabors allowed us to quantify the difference between the successive targets’ orientations. To track any bias on a trial-to-trial basis, we used a reproduction task, where participants replicated the target orientations at the end of the trial. The Lag (the number of items T2 appeared after T1) was varied to measure the time course of attentional selection. We characterised accuracy for correctly reproducing the target orientation (within ±30 degrees of the presented orientation) separately for the two targets.

To provide an initial test of the model’s prediction, we first split trials depending on the degree of difference in orientation between the targets (Figure 2C). T1 accuracy was significantly higher when the orientations of T1 and T2 were similar than when they were different (Figure 2C). This effect was more pronounced when the targets were presented sequentially at Lag 1. A 2 (Similarity; Similar, Dissimilar) x 5 (Lag; 1,2,3,5,7) repeated-measures ANOVA confirmed a significant effect of Similarity *(F*(1,21) = 42.08, *p* < 0.001,*η*^*2*^ = 0.23), Lag *(F*(2.64,55.38) = 29.22, *p* < 0.001,*η*^*2*^ = 0.24), and the interaction between these factors *(F*(2.43,51.00) = 10.20, *p* < 0.001,*η*^*2*^ = 0.08) on T1 accuracy. For T2 accuracy (calculated only on trials where T1 was correct), targets with dissimilar orientations showed the classic AB phenomenon, with significantly lower accuracy at Lags 2 and 3 which recovered by Lags 5 and 7.

**Figure 2.**
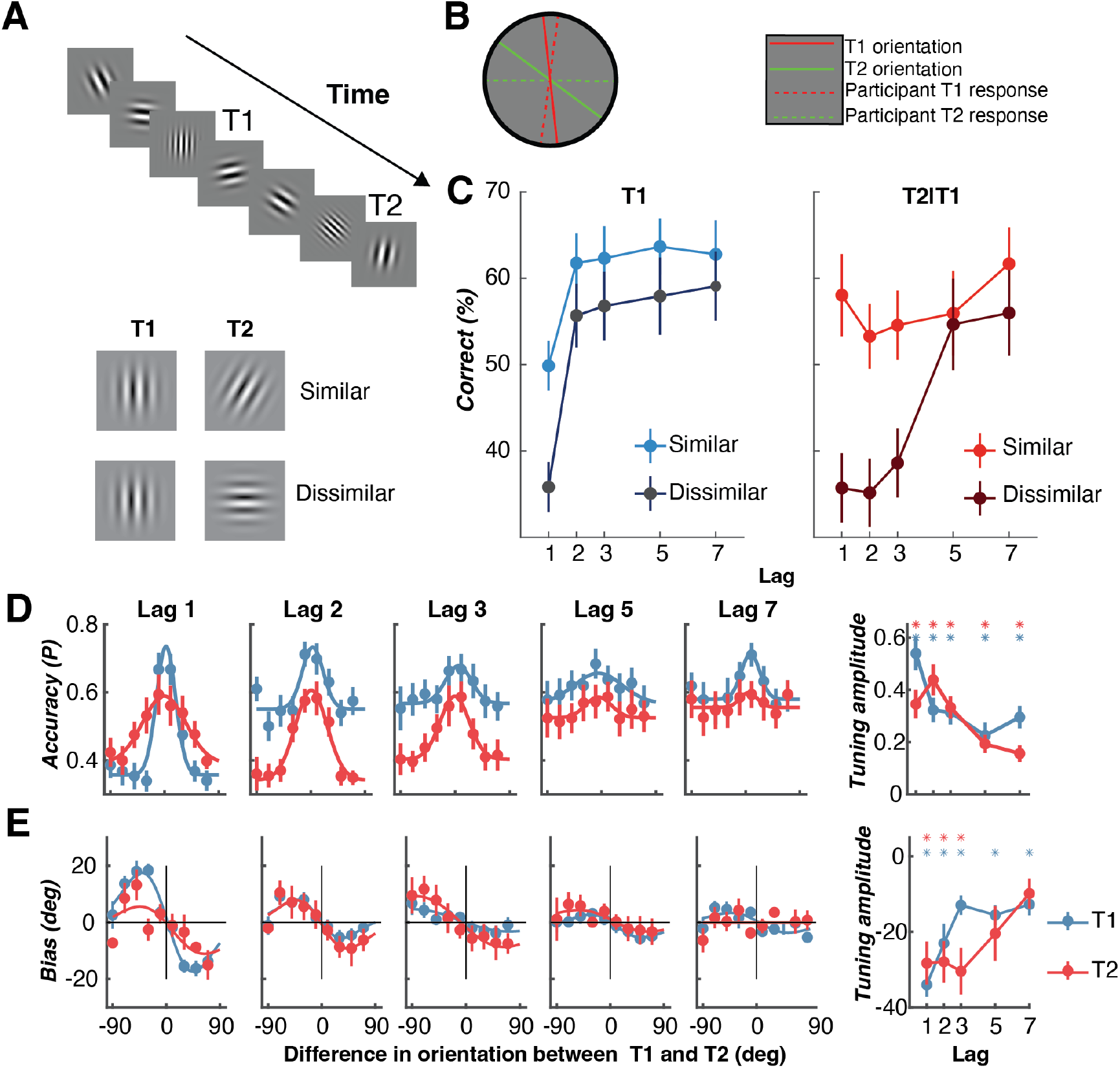
Using a Gabor RSVP task to determine the effect of similarity on temporal target selection in Experiments 1 and 2. (A) Schematic of the Gabor RSVP task. Participants viewed a stream of Gabors and were required to reproduce (B) the orientation of the two higher-spatial frequency targets. Trials where the participant reproduced the orientation within 30° of the true orientation were classified as correct. Lag is defined as the difference in item positions between T1 and T2. Trials were classified as similar or dissimilar depending on the orientation difference between the two targets. (C) Behavioural accuracy for T1 and T2 for each Lag for Experiment 1. (D) Accuracy as a function of the difference in orientation between T1 and T2. For T2, only trials where T1 was correctly reported were included. Dots are the psychophysical data, and the fitted line is the Gaussian function used to quantify the tuning. The right-most panel shows the amplitude of the Gaussian, fitted to each participant’s data separately, for all lags. The asterisks show when the amplitude was significantly greater than 0 (Bonferroni-corrected). (E) The bias in reported orientation by the difference in orientation between T1 and T2. Data have been replotted from Tang, et al.^19^. The fitted line is the first derivative of a Gaussian used to quantify the bias. Across all panels the error bars and shading indicate ±1 standard error.

Notably, however, the AB was significantly reduced when the successive targets had similar orientations, with participants being able to report both items regardless of the interval. A 2 (Similarity) x 5 (Lag) repeated-measures ANOVA supported these observations. There were significant effects of Similarity *(F*(1,21) = 6.76, *p* = 0.01, *η*_*p*_^*2*^ = 0.24), Lag *(F*(2.29,48.08) = 9.56, *p* < 0.001, *η*_*p*_^*2*^ = 0.31), and the interaction between these factors *(F*(3.11,65.32) = 8.31, *p* < 0.001, *η*_*p*_^*2*^ = 0.28) on T2 accuracy. To confirm the difference between conditions, separate repeated-measures ANOVAs for each Similarity condition showed a significant effect of Lag for dissimilar *(F*(2.39,50.23) = 13.82, *p* < 0.001, *η*_*p*_^*2*^ = 0.41), but not similar *(F*(3.10, 65.15) = 1.89, *p* = 0.12, *η*_*p*_^*2*^ = 0.08), trials.

The effect of target similarity on accuracy extends beyond a simple split between similar and dissimilar trials. We re-analysed the data from Experiment 1 but now plotted accuracy as a function of the difference in orientation between the two targets (Figure 2D). This revealed an orientation-selective effect of similarity on accuracy for both T1 and T2 with the same tuning predicted by our model. The highest accuracy was observed when the target orientations were most similar, and this decreased as the orientations of the two targets became more dissimilar. Remarkably, when the targets were similar, T2 accuracy was comparable to overall T1 accuracy across all Lags, suggesting the AB was not present for similar targets. To further quantify this tuning, Gaussian functions were fit to each participant’s accuracy for each Lag (see Methods). This showed significant orientation modulation of accuracy for all targets and lags. A 2 (Target; T1, T2) x 5 (Lag; 1,2,3,5,7) within-subjects repeated-measures ANOVA showed the magnitude of the tuning decreased with Lag *(F*(4,84) = 6.84, *p* < 0.001,*η*_*p*_^*2*^ = 0.25), with the decrease in Lag being dependent on the interaction with Target (*F*(4,84) = 3.10, *p* = 0.02, *η*_*p*_^*2*^ = 0.13), but with no main effect of Target (*F*(1,21)<1).

The finding that accuracy is determined by the orientation difference between successive targets extends our previously-reported effect where the perceived orientations of targets are attracted to each other^20^. Figure 2E shows bias in reported orientation by the orientation difference between the targets. While on average the orientation errors are centred around 0°, there are systematic biases in perceived orientation that emerge depending on the orientation difference between the targets. When T1 is clockwise of T2, the perceived orientation of T2 is biased clockwise for positive differences and the corresponding effect when T1 is anti-clockwise of T2. These effects are directly analogous to serial dependency effects where the largest attraction occurs when the targets are ~45° apart^28,29^. This is true despite the fact that unlike most studies examining these effects, the targets in our task were presented much closer together in time and perceptual judgements were made after both stimuli had been presented. Similar to accuracy, a 2 (Target) x 5 (Lag) within-subjects repeated-measures ANOVA revealed that the magnitude of the bias decreased across Lag *(F*(3.00,63.10) = 8.72, *p* < 0.001, *η*_*p*_^*2*^ = 0.29), with the decrease in Lag dependent on the interaction with Target (*F*(3.00,63.06) = 4.22, *p* = 0.004, *η*_*p*_^*2*^ = 0.17), but with no main effect of Target (*F*(1,21) = 2.35, *p* = 0.14, *η*_*p*_^*2*^ = 0.10).

Figure 3 uses the model to predict the exact pattern of behavioural results observed in Experiment 1. We first binned the trials into the orientation difference between T1 and T2 for accuracy at the height of the AB (Lag 2, Figure 2A, red points). We then plotted the amplitude of the orientation channels (given by the vector sum) to these data for all differences in orientation between T1 and T2 (solid line). The figure shows that the model provides an excellent fit for the behavioural results with the magnitude of the population response showing the same orientation dependency. Next, we used the same trials but found the model’s prediction of the perceived orientation (given by the vector mean). There is a strong theoretical relationship between the population response and behavioural accuracy. Numerous previous studies have found that accuracy is higher when there is a larger neural response. Furthermore, global workspace theories predict that conscious awareness of targets only occurs when the neural response exceeds a certain threshold^24,25^. Our findings from Experiment 1 fit well with this prediction as T2 is more likely to be missed when there is a large orientation difference and a predicted smaller neural response.

**Figure 3.**
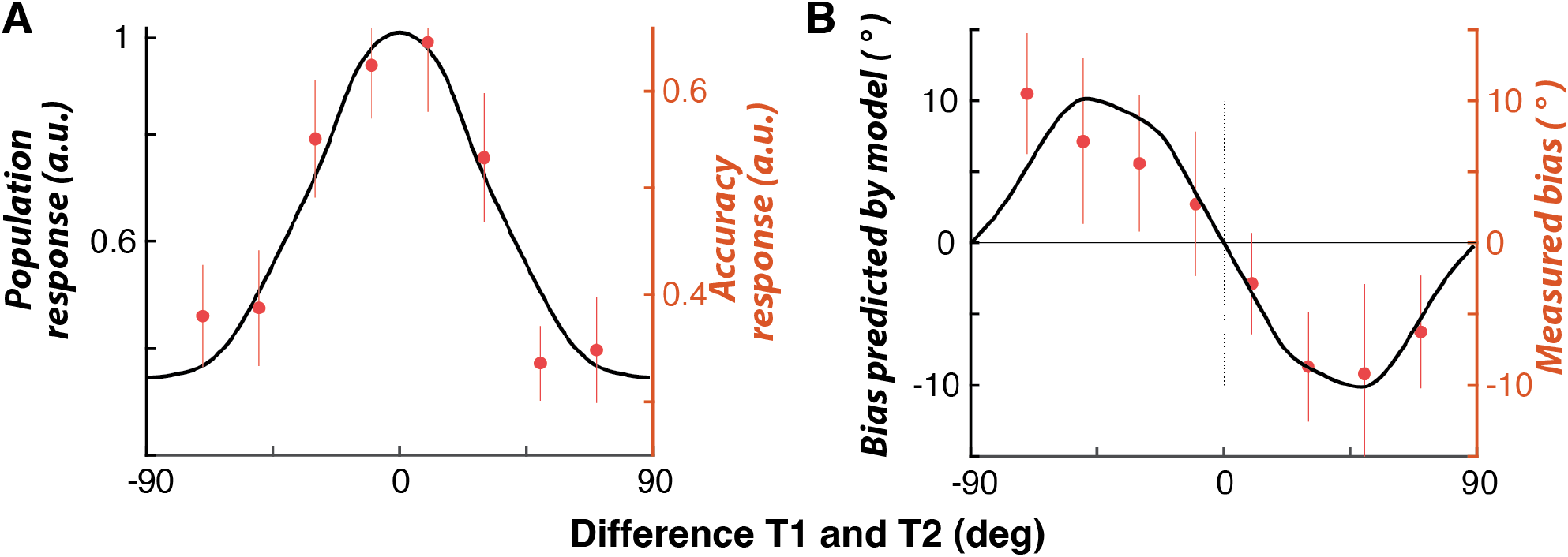
(A) The model’s predictions (black line) against behavioural data (red points) from Experiment 1, replotted from Figure 1. Bias in reporting orientation. (B). Same as C but with accuracy for T2 Lag 2 (for T1 correct trials). Error bars indicate ±1 standard error.

### Experiment 2 – Testing the representational model of the AB against EEG activity

To investigate the time course of these effects across different brain areas, we examined EEG activity from a second group of participants (N = 23) who completed the same dual-target Gabor RSVP task as used in Experiment 1. To increase experimental power, we only presented targets at Lags 3 and 7, but all other details of the experiment were the same. We first investigated the time course and neural loci of these representation effects on attention by examining the event-related potentials (ERPs; Figure 4A). The similarity between the targets caused two distinct patterns; one early sensory response and a later frontal effect (Supplementary Figure 1). Shortly after T2 presentation (203 to 257 ms), there was a larger response to dissimilar relative to similar stimuli in clusters of both visuo-parietal and frontal electrodes. The effect was strongest over the occipital sensors, with the sign of the activity flipped in the frontal sensors suggesting a common EEG dipole located in the visuo-parietal areas. Furthermore, the time course suggests the effect originates in sensory processing areas, in which stimulus-driven activity is evoked rapidly. This was followed by a later effect (545 to 594 ms) in frontal electrodes, where again dissimilar trials resulted in a larger response compared with similar trials. Interestingly, the AB is commonly associated with late-stage ERP differences (P300, ~300-500 ms) following the second target^18^. We have also recently shown very early effects (~100-200 ms) using this task^20^, which is consistent with the time course of other work examining the difference between consciously- and unconsciously-perceived stimuli^30^ and work showing the AB is related to changes in processing in the primary visual cortex (V1)^19^. The differences between the latency may reflect the different processing level for the stimuli (Gabors versus letters). These results point to the topography and timing (Supplementary Figure 1) of the changes in occipital activity, with the earliest difference likely reflecting a filter/template emerging early in the visual processing hierarchy following T1 presentation.

**Figure 4.**
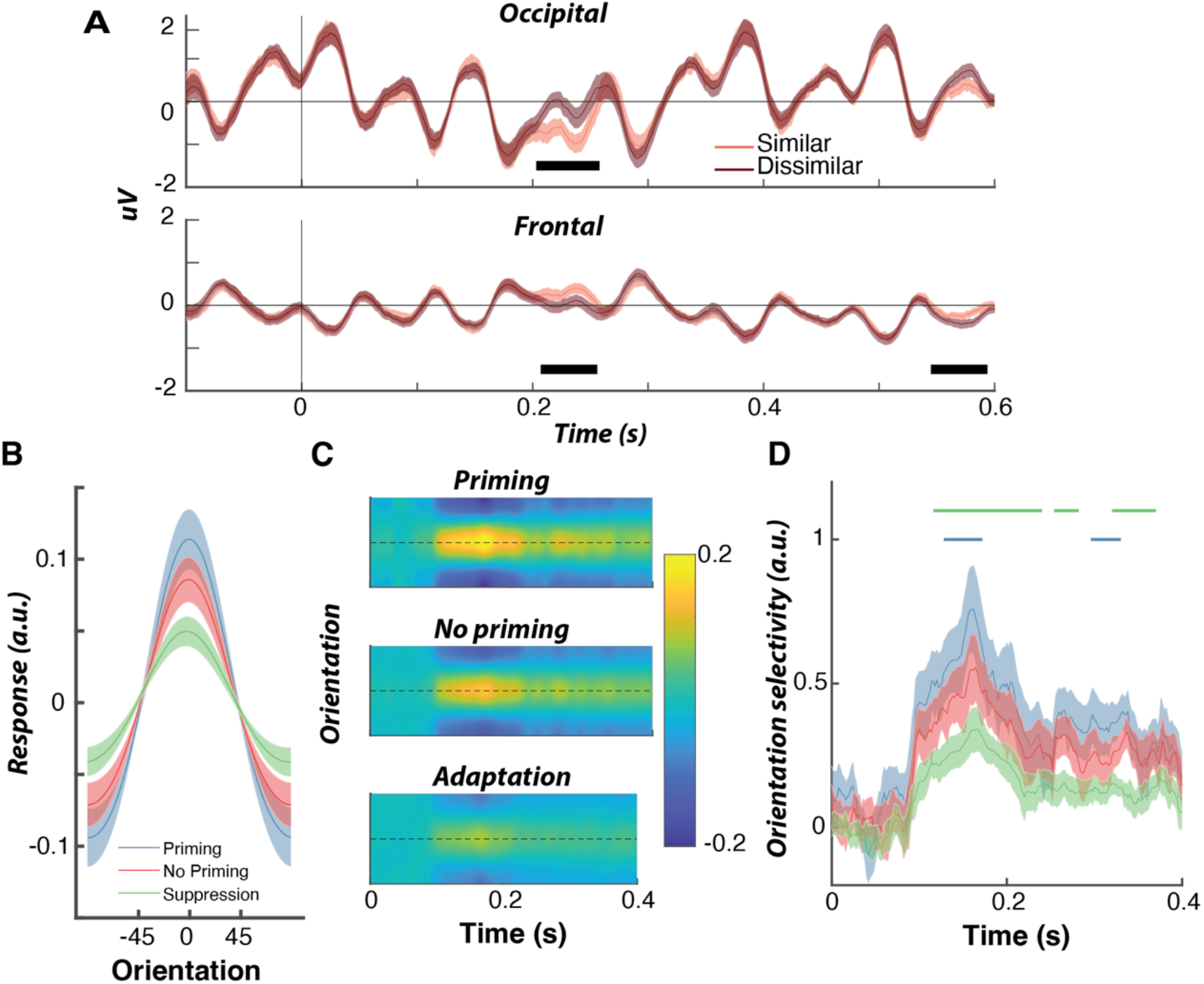
(A) The event-related potentials (ERP) for Experiment 2. ERP signals are averaged across the cluster-permutation significant occipital-parietal electrodes. The shaded area indicates cluster-corrected significant differences between the two conditions (similar and dissimilar). (B) The response of the population of neurons to different T2 orientations relative to T1. Forward encoding of orientation selectivity in EEG activity from Experiment 2 with different models used to predict orientation response. Results are averaged between 100 and 250 ms after the stimulus. (C) Time-resolved forward encoding. (D) Gaussians were fit to the time-resolved encoding results to quantify orientation selectivity with greater Gaussian amplitudes reflecting greater selectivity. Horizontal lines indicate significant cluster-permutated differences between Priming and No priming (blue) and Suppression and No-priming. Across all panels the error bars and shading indicate ±1 standard error.

To provide a further test of the representational model, we used its outputs to predict the pattern of EEG activity recorded in Experiment 2. To do this, we used a modified version of forward encoding modelling, which uses a multivariate regression-based approach to find patterns of neural activity selective for feature-based activity, in this case orientation selectivity^20,23,31–33^. As this is a regression-based approach, it gives the opportunity to test different models of selectivity to see which most accurately predicts EEG activity on a trial-by-trial basis. We used three versions of the model to generate different regressors to predict the orientation of T2. In the first model, T1 was assumed to exert no influence on the response to T2 (no priming), with the regressor values being the same as the initial unbiased model (Figure 2A). The second model used the T1-priming model, where the channels selective for T1 have increased sensitivity relative to non-selective channels, to predict the response to T2 (Figure 4C). The third model was the classic adaptation model in which T1 decreased the sensitivity of the activated channels. The third model was included for two reasons. First, one theory argues that capacity limitations of channels determine the magnitude of the AB, with the largest AB occurring when the stimuli activate the same channels^34^. T1 would thus decrease the sensitivity of the activated channels in an adaptation-like fashion. Second, this model provides an important test to determine whether using the identity of T1 to modulate the activation pattern associated with T2 will necessarily lead to better multivariate decoding.

This analysis revealed strong orientation selectivity for all three models (Figure F), with orientation presented at T2 being decoded significantly above chance. We fit Gaussian functions to each participant’s encoding results at each time point to quantify the time-resolved orientation selectivity (Figure 4D). This showed that the priming model led to significantly better encoding of orientation selectivity, emerging ~100ms after stimulus presentation. The adaptation model, on the other hand, led to significantly worse decoding than the no-priming model. Overall, the model for orientation processing in which T1 primes the system for stimuli which are represented similarly captures the behavioural data well. As a bonus, this model explains more of the EEG activity than can be explained without this change in stimulus selectivity.

### Experiment 3 – Similarity effects extend to alpha-numeric digit targets

Experiments 1 and 2 provide support for our unifying model of temporal attention, in which attention decreases sensitivity to items that have dissimilar representations to the target. Our results are consistent with the idea that attending to an initial target creates a transient filter that determines whether a second target will be consciously accessible. Priming the attended stimulus representation allows only similarly-represented stimuli to be accurately reported.

We next examined whether this representational model of the AB extends beyond the simple orientation space into more complex feature space. To do this, we asked whether in a traditional RSVP stream consisting of alphanumeric items^3,5^, the AB is reduced when the targets are closely-spaced numbers. We chose an alphanumeric RSVP task because numerical representation follows a mental ‘number line’, with numbers increasing in magnitude from left to right along the line. This numerical representation has been demonstrated at both the cognitive and neural levels^35–37^. Furthermore, alphanumeric RSVP tasks are commonly used to measure the AB.

To test whether these similarity effects extend to numerical processing, we used a dual-target RSVP task in which the targets were numbers and the distractors were letters (Figure 5A^38^). Participants were required to report the targets using a keyboard. To determine whether target similarity affects conscious awareness, we calculated reporting accuracy by the numerical difference between the two targets (Figure 5B). Consistent with our model’s predictions, participants’ ability to report T2 depended on its similarity with T1, when the targets were separated by short (Lag 2) intervals, whereas this did not occur at longer intervals (Lag 7). Accuracy was maximal when the targets were only separated by one digit and worse at larger differences in magnitude, exhibiting the tuning-like effect seen in Experiment 1, but this time for numbers. Unlike the previous task, the targets in this experiment never had the same physical identity, which likely reduced the magnitude of the similarity effect. A 2 (Target; T1, T2) x 8 (Difference; −4 to +4) ANOVA on Lag 2 data confirmed that accuracy was higher for T1 than T2 (*F*(1,42) = 149.99, *p* < 0.001, *η*_*p*_^*2*^ = 0.78) and the effect of Difference only occurred as an interaction with Target (*F*(5.68,238.70) = 2.32, *p* = 0.04, *η*_*p*_^*2*^ = 0.05), while there was no main effect of the Difference (*F*(5.63,236.34) = 1.45, *p* = 0.20, *η*_*p*_^*2*^ = 0.03). For Lag 7 trials, accuracy was only higher for T1 than T2 (*F*(1,43) = 16.87, *p* < 0.001, *η*_*p*_^*2*^ = 0.28), while there was no significant interaction or main effect of Difference (*Fs* < 1). Overall, T1 appears to constrain conscious awareness of T2 based on high-level, conceptual representations.

**Figure 5.**
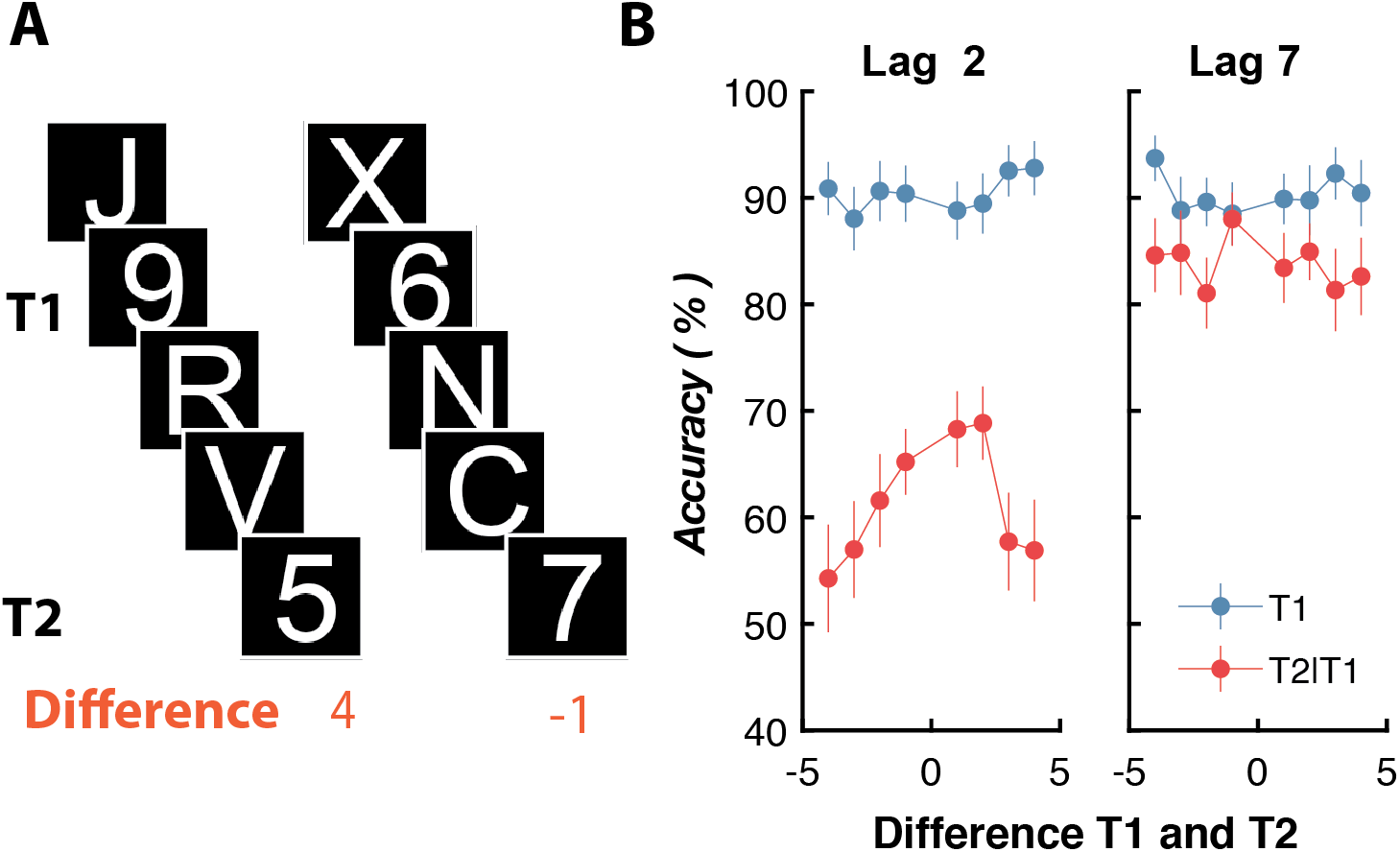
Similarity effects for target perception extend to more complex alphanumeric representations in Experiment 3 (N = 49). Targets were numbers and distractors were letters. In one trial, the identity of the target did not repeat, so targets were never the same number. The difference between targets was found for each trial. (B) Accuracy for Lag 2 and 7 as a function of the difference between targets. The error bars indicate ±1 standard error. Data were originally published in Tang, et al. ^33^.

### Experiment 4 – Target priming in alphanumeric letters is driven by representational similarity

So far, we have shown that detecting a target causes the brain to create a temporary filter which allows for selection of targets items with similar neural representations. We have also shown that similarity can be defined by intrinsic, low-level features such as orientation as well as high-level representations like numerical value, which are known to have a relatively simple representation of sequential order. Our next goal was to extend our findings to alphabetical letter targets, which have been widely used to measure attentional selection in humans^5,39^. Letters have a very complex representational structure^40,41^, which is determined from low-level visual information, such as object form and orientation to high-level information that is learnt through experience, for instance the order in which they commonly appear in language and through their alphabetical order. To examine how alphabetical representation affects the AB, we used a similar task as in the previous experiments, but now presented dual-target RSVPs (Figure 6A) with letter targets and number or letter-like distractors^42,43^. The participants (N = 119) were required to type the presented target letters at the end of the presentation (see Methods for details). Here we defined representational similarity as the difference in the alphabetical order between the target letters. Seminal work suggests English speakers have a representation of alphabetical order that is analogous to the number line representation explored in Experiment 3^44–46^. These studies found participants are significantly faster judging the relative order of two letters when they are close than when they are further apart in the alphabet.

**Figure 6.**
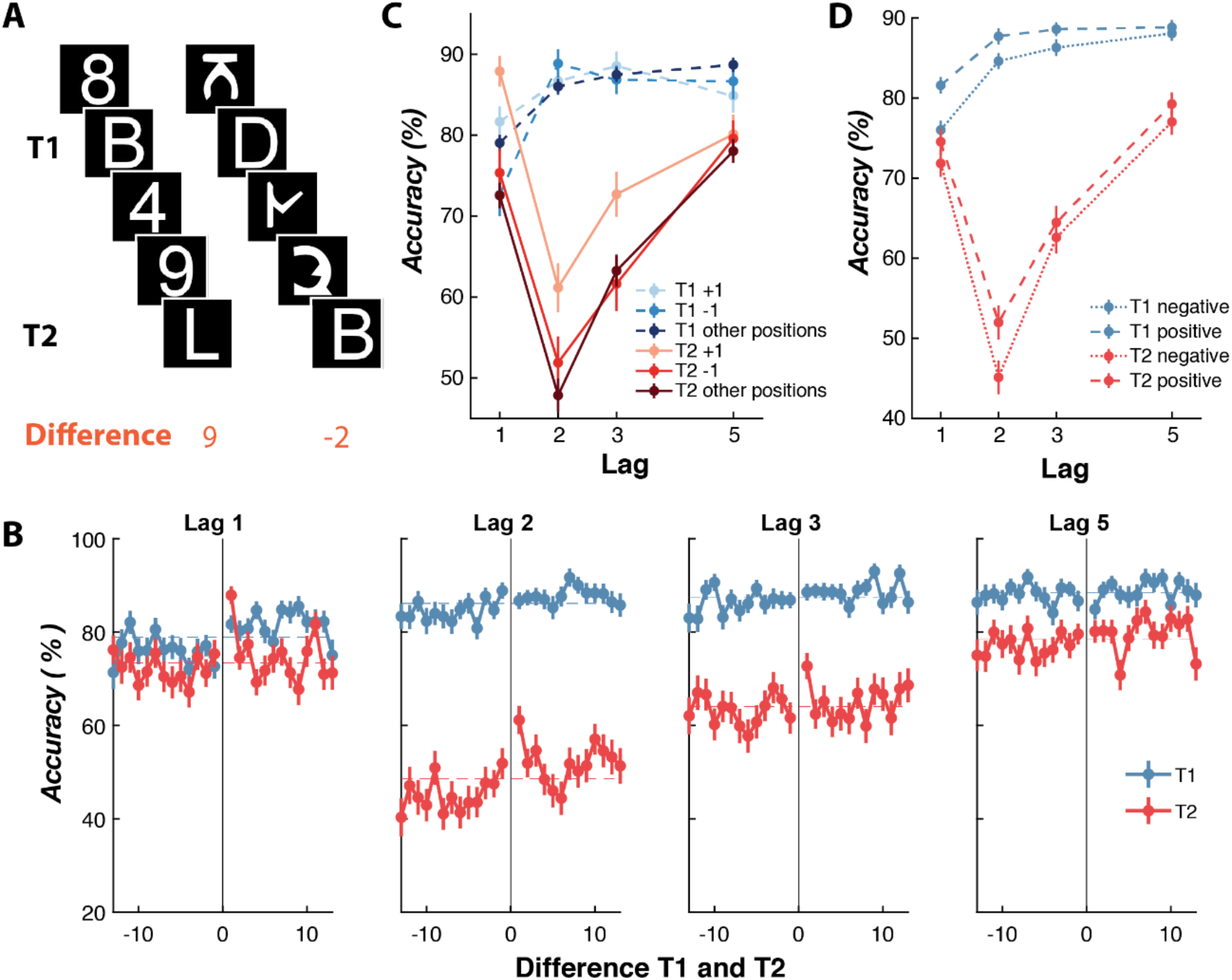
Target similarity for letter targets defined by the difference in alphabetical order in Experiment 4 (N = 119). (A) Targets were letters and distractors were either digits or pseudo-letters which had a high structural similarity with letters. (B) Accuracy by the alphabetical difference between letters. The dashed horizontal line indicates the mean accuracy for all differences. (C) Accuracy for +1 and −1 difference relative to all other differences between T1 and T2. (D) Switching of perceived target order by the difference between T1 and T2 for all lags. Across all panels the error bars indicate ±1 standard error.

To quantify this effect, we compared the accuracy by Lag for +1,-1 and all other differences using a 3 (Difference; +1,-1, other positions) x 4 (Lag; 1,2,3,5) repeated-measures ANOVA, separately for T1 and T2. For T2, this revealed accuracy was affected by the Difference (*F*(1.50,176.83) = 23.12, *p* < 0.001, *η*_*p*_^*2*^ = 0.16), the Lag (*F*(2.76,326.27) = 74.63, *p*<0.001, *η*_*p*_^*2*^ = 0.39) and the interaction between these factors (*F*(4.50,531.32)=3.36, *p* = 0.007, *η*_*p*_^*2*^ = 0.03). Post-hoc tests with a Bonferroni correction showed that the +1 difference condition had higher accuracy than −1 and the others (*all ps* < 0.001, whereas there was no difference between −1 and the others (*p* = 0.34). For T1, there was no overall effect of the Difference (*F*(1.80,212.78) = 1.74, *p* = 0.18, *η*_*p*_^*2*^ = 0.02), whereas Lag (*F*(2.86,337.81) = 25.97, *p* < 0.001, *η*_*p*_^*2*^ = 0.18) and the interaction between these factors (*F*(4.74,558.88) = 3.33, *p* = 0.007, *η*_*p*_^*2*^ = 0.03) significantly affected accuracy. The interaction is driven by the lower accuracy for −1 Difference at Lag 1 relative to the +1 and others (Bonferroni *p*=0.002).

Another aspect of these data worth noting is that accuracy in general was lower when there was a negative difference (e.g., D followed by B) relative to a positive difference (e.g., B followed by D). This is possibly because we more commonly rehearse the alphabet in one direction (A through to Z) and rarely in reverse (Z through to A). To quantify these effects, we compared accuracy for all negative and positive differences (Figure 6D). A 2 (Difference; Positive and Negative) x 4 (Lag; 1,2,3,5) repeated-measure ANOVA was conducted for each target accuracy. For T1, accuracy was affected by the Difference (*F*(1,118) = 57.38, *p* < 0.001, *η*_*p*_^*2*^ = 0.33), Lag (*F*(2.72,321.47) = 82.10, *p* < 0.001, *η*_*p*_^*2*^ = 0.41), and interaction between these factors (*F*(2.84,335.22) = 5.69, *p* = 0.001, *η*_*p*_^*2*^ = 0.05). Follow-up post hoc tests indicated that the interaction arose because accuracy was significantly higher for Lags 1 and 2 (*all Bonferroni ps* < 0.01), but not Lags 3 and 5 (*all ps* > 0.15), for Positive relative to Negative differences. For T2, accuracy was affected by the Difference (*F*(1,118) = 35.50, *p* < 0.001, *η*_*p*_^*2*^ = 0.23), Lag (*F*(2.42,285.36) = 180.82, *p* < 0.001, *η*_*p*_^*2*^ = 0.60), and interaction between these factors (*F*(2.78,328.33) = 4.18, *p* = 0.006, *η*_*p*_^*2*^ = 0.03). The significant interaction arose because accuracy was significantly higher for Lag 2 (*Bonferroni p* < 0.001), but not the other Lags (*all ps* > 0.5), for Positive relative to Negative differences. This suggests the representational similarity effect is largest for the Lag at the maximal depth of the AB (Lag 2), and aligns with the results from Experiment 1 with Gabors.

### Experiment 5 - Representational similarity from mouse visual cortical neuronal activity and a Deep-neural network both predict human behavioural performance

Alphanumeric letters are a powerful stimulus set for probing visual representations because they encompass multiple levels of abstraction; from low level properties such as line orientation and curvature, to high level statistical relationships which are learnt through experience with the language, such as the frequency with which two letters commonly appear together (e.g., ‘th’ is the most common pair in English). Using letters, we have so far explored how a high-level feature, alphabetical order, affects the ability to bring multiple items into conscious awareness. In Experiment 5, our aim was to characterise the different contributions of low- and high-level features on the representational account of the AB.

To determine how the low-level features of letters affect the AB, we examined representational similarity in population activity recorded from mouse (C57BL) visual cortical neurons. Mice are increasingly the dominant species used in visual neuroscience, and, like primates, have multiple visual cortical areas which respond to increasingly complex information. Furthermore, the majority of neurons in the primary visual cortex are selective for combinations of orientation, retinotopic location and spatial frequency^47–49^.

Most importantly, representational similarity in the population response should only be due to low-level features because mice have limited (or no) exposure to visual letters.

First, we asked whether the activity evoked by letters in a large population of neurons could be used to predict the human AB from Experiment 4 (Figure 7A). Awake mice passively viewed the stimuli which were, as in Experiment 4, alphanumeric English letters presented in RSVP sequences at 10 Hz (Figure 7B; see Methods for details). We recorded activity in primary visual cortex using Neuropixel electrodes over 7 sessions in three mice (2510 units recorded). Many of the individual units showed strong selectivity for certain letters. The representative neuron shown in Figure 7C responded strongly to W, C, D, K, which all contain oblique orientations or curved features, whereas spiking was suppressed (relative to baseline) for T, V, and F. Across the entire population of neurons significant stimulus selectivity emerged shortly after stimulus presentation (Figure 7D). We trained a linear multivariate decoder to discriminate the identity of the letters presented in each trial from the population response recorded in each session. This decoder successfully predicted the presented letter on previously unused test trials, with significant decoding emerging around 150 ms after stimulus onset. These results reveal that letters drive unique patterns of activity at the population level in mouse primary visual cortex.

**Figure 7.**
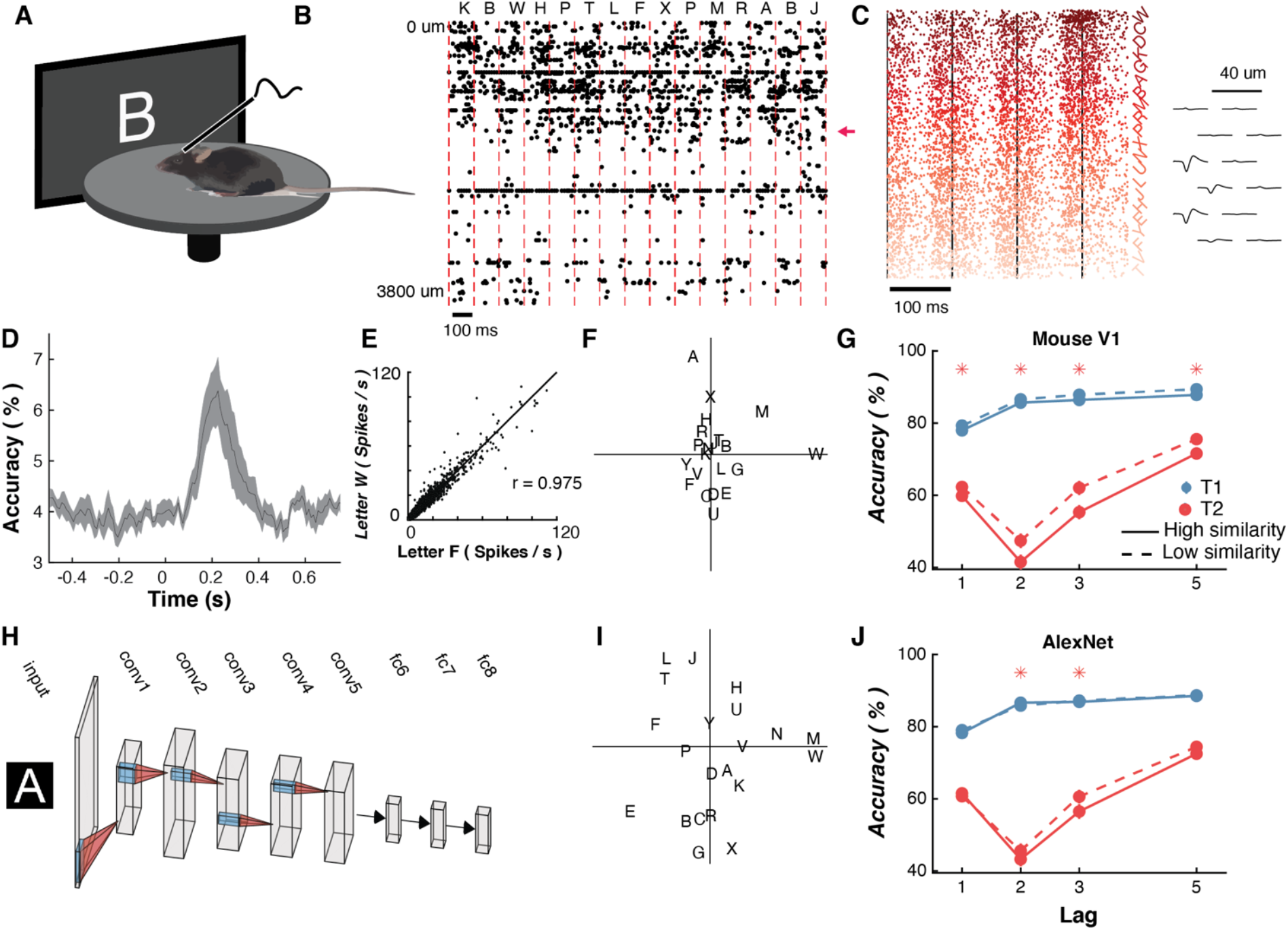
Low-level representational similarity from mouse primary visual cortex neurons and a biologically-inspired Deep Neural Network (AlexNet) predict human conscious awareness in Experiment 4. (A) To determine representation similarity of letters in neuronal activity, awake head-fixed mice (N = 3) passively viewed 10 Hz RSVP streams consisting of the same target letters as those used in Experiment 4 while neuronal activity was recorded using high-density extracellular electrodes (Neuropixel, N = 2510 units across 7 session). (B) Spiking activity of all simultaneously recorded neurons (N = 512) in one of 7 sessions to the RSVP. Each line is one neuron, and each dot is a spike. The more superficial neurons show consistent evoked responses following presentation of the letters. (C) An example neuron’s response to all presentations of the letters across the experiment. The position of this neuron in B is shown by the red arrow. Each row is a presentation. The trials have been sorted by the most responsive letters for this neuron. The neuron shows maximum responses to W, B, and C, while showing suppression relative to baseline spiking activity, to T and V, which emerges most strongly ~180 ms after stimulus presentation. The unit’s waveform across 12 of the tightly spaced channels is shown in the insert. (D) Multivariate decoding of letter identity from population neuronal activity. (E) An example of representational similarity analysis for two letters. The responses of all recorded neurons (dots) in primary visual cortex evoked by letters W and S. (F) The representational similarity analysis (RSA) results for mouse V1 representations shown using multi-dimensional scaling. Letters that have more similar representations are plotted more closely to one another. (G) Asterisks show Bonferroni-corrected t-test differences between similarity conditions, calculated separately for T1 and T2. (H) The architecture of AlexNet. Each box is a separate layer. (I) RSA results for AlexNet (Conv2 layer) plotted using multi-dimensional scaling. (J) Same as G, but based on AlexNet (Conv2) representations. Across all panels the error bars and shading indicate ±1 standard error.

As the previous analysis confirmed letters are represented uniquely in mouse visual cortex, we next determined the representational similarity of the letters in mouse primary cortical activity. To do this, we performed a representation similarity analysis (RSA^50,51^) on the population responses of the recorded neurons. As expected, because mouse V1 neurons are selectively responsive to both orientation and retinotopic location, and have had limited (or no) prior exposure to English characters, similarity mainly appears to be driven by low-level factors such as orientation and curvature (Figure 7F). For example, P, R and H were all represented similarly, perhaps by virtue of possessing a left vertical line-segment and a horizontal line-segment at their midpoint. Likewise, O and D share basic geometric features and were represented similarly. To determine whether the low-level similarity between letters affected human observers’ ability to detect two stimuli presented in rapid sequence in Experiment 4, we examined whether the mouse similarity representation of the letters could be used to predict human behavioural performance (Figure 7G). To do this, we used a median split on the mouse V1 population data to classify the representation similarities of each pair of letters into high- and low-similarity pairs. We then used this to categorise each trial of the results in Experiment 4 as ‘high’ and ‘low’ similarity (Figure 7G). T2 performance was significantly higher when it had a high similarity with T1 for all Lags. T1 performance was relatively unaffected by its match with T2. How similarly the letters were represented in patterns of neuronal activity in mouse primary visual cortex therefore is related to the ability of humans to detect targets. This finding suggests that the low-level similarity of the targets, in addition to the learnt representations, strongly influence whether the second target reaches conscious awareness.

We next used a deep neural network (DNN; AlexNet), which has been trained for image labelling of natural scenes, to determine low-level similarity in human participants^52^. The network (Figure 7H) is based on a processing hierarchy similar to that of the primate visual system^53^ and has been shown to predict the response of individual neurons in monkey area V4^54^ in addition to visual responses in human fMRI and EEG activity^55^. We then performed an RSA for each layer by correlating the population response for all the letters (Figure 7I, Supplementary Figure 2). To do this, we found the activation for each level of the network for the 21 letters we presented to the participants in Experiment 4. This analysis revealed that low-level structural features largely drive the representational similarity, especially for the earlier layers (Supplementary Figure 2). For layer Conv2, letters with oblique lines (X,V,A) had similar representations, as did the letters with curves (B,G,D). The low-level, feature-based aspects of the stimuli quantified by the DNN also affected performance. Like mouse V1 representations, T2 was more likely to be reported when T1 was represented similarly in AlexNet (Figure 7J). Overall, these results again suggest that human observers’ ability to bring the second of two rapidly presented stimuli to consciousness depends on its representational similarity to the first item.

Finally, we aimed to unify our initial results showing low-level structural features, with high-level learnt factors, and determine the relative influence of each factor on whether targets reach the level of conscious awareness. To do this, we used a regression-based approach to determine the amount of variance explained by different high- and low-level representational factors. We correlated the human performance data in Experiment 4 with representational similarity, defined by Mouse V1 neuronal activity, AlexNet (Conv2), letter order and frequency of letter co-occurrence (Figure 8). Co-occurrence of letters (or bigrams) captures the probability that any two letters commonly occur together, and is measured from the Google Books One Million Book corpus analysis. We used this as a high-level factor that is superordinate over letters, as participants often report that they can detect both targets if they regularly co-occur (e.g., in NY, AM) and everyday exposure to language is expected to affect our representation in a similar manner to alphabetical order.

**Figure 8.**
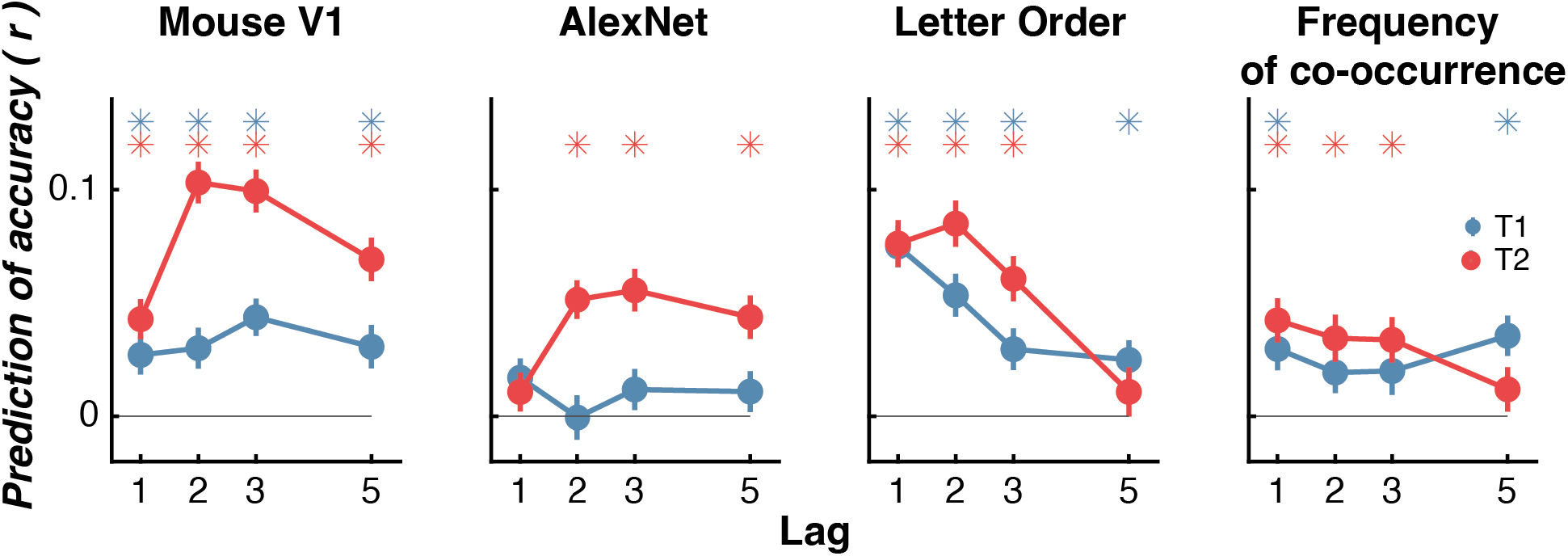
The effects of intrinsic low-level and learnt high-level representational similarity on access to conscious awareness for Experiment 4 and 5. Asterisks show Bonferroni-corrected one-sample t-test showing correlation is greater than 0, calculated separately for each target. Across all panels the error bars indicate ±1 standard error.

Like the previous results, this final analysis revealed that representations derived from mouse V1 neuronal activity best predicted whether participants could report T2. The letter order (Figure 8) was the next best predictor of T2 activity, again with the best performance during the depth of the AB. Similarly, if T2 frequently follows T1, participants were more likely to correctly report T2 for Lags 1, 2 and 3. Interestingly, T1 only showed a significant effect of T2 for Lags 1 and 5, those where T2 had the highest accuracy. Overall, it appears that both low-level intrinsic aspects and high-level learnt statistical relationships combine to determine neural representations of letters. When we see a target, it activates the representation which creates a temporal filter, allowing only similarly represented items to channel through but suppressing dissimilar items.

## Discussion

One of the most consistently cited truisms in the neuroscientific and psychological literature is that limitations in conscious awareness occur when stimulus demands exceed the processing capacity of the brain. This belief is based on numerous demonstrations showing acute failures in awareness when multiple, competing demands occur^56,57^. The current results point to a new interpretation of one of the most widely cited of these perceptual limitations, the attentional blink (AB)^3–5^. Here we developed and tested a new unified model of temporal attention with several lines of converging evidence showing that the act of attending to one item causes automatic filtering of subsequent items with dissimilar representations. This is because the representation of the first target becomes enhanced relative to the unattended representations. The ability to bring the second item to conscious awareness depends strongly on its representational similarity with the first target. If the targets are represented in a similar manner, both can be reported whereas the second is suppressed from conscious awareness if it is represented differently to the first. This filtering is the cause of the AB and unifies it with spatial and feature-based attention.

Experiments 1 and 2 showed this with simple, low-dimensional Gabor stimuli that have a well-characterised representational space which is easily quantifiable^32^. The EEG results showed similarity affects very early stages (~200-250 ms) of processing in visual-parietal areas. This very early effect of similarity suggests that this may reflect a filter that was existing prior to T2 being presented and is due to the ongoing processing of T1. We found that the orientation-selective bias and accuracy results can be explained by a model where the orientation of T1 suppresses the non-activated representations. This model was better at predicting the EEG activity recorded during the task than models where T1 does not affect the response to T2 or one that suppresses activity of the activated channels.

Experiment 3 extended these findings to the representation of alpha-numeric numbers which differ in their position on the number line rather than low-level structural features of the stimuli. Consistent with the initial results, this experiment found that T2 was less likely to be missed if it had a similar number to the first (i.e., closer on the number line). The final experiment showed that T2 was more likely to be detected if it had a similar representation to T1 based on high-level learnt aspects (letter order, frequency) or low-level features of the letters. Our results suggest that detection of a target causes the brain to establish an automatic filter which results in selection of items matching the identity of the leading items and suppression from consciousness for those items which do not match.

The current work leads us to re-evaluate the most widely studied task used to interrogate human limitations in temporal awareness; the attentional blink (AB). This refers to the profound inability to detect the second of two targets when it follows within 200-500 ms of the first^3,5^, and for which there have been numerous theories, mostly arguing that the second target deficit is caused by a lack of processing resources for consolidating targets into working memory. The current results instead point to an early sensory driven filter based on representational similarity. In some ways this idea builds on the over-investment account of the AB, where detecting T1 causes an over-allocation of attentional resources to the RSVP causing distractors to be selected. Our theory that attending T1 causes the brain to develop an involuntary filter based on the activated representation is not inherently based on resource limitations. In many ways this idea is consistent with a Bayesian predictive coding account of perception^32,33,58,59^. T1 establishes a ‘prior’. This idea fits with everyday perception, sensory input generally changes slowly^60^ most items that are drastically different to what came before should likely be discarded as noise.

A number of previous studies have examined the effect of target similarity on the magnitude of the AB. Consistent with the current results, the magnitude of the AB in those studies was reduced when the targets had the same features, but different orientations, compared to when they have different features^61,62^. The current results are consistent with the priming literature, where repetitions tend to lead to perceptual facilitation^63^It has also recently been reported that natural images that are similar to one another, with similarity quantified using a DNN approach as in the current work, lead to a smaller AB^51^. But there are other studies, referred to collectively as repetition blindness studies, that seem to suggest that similarity of targets leads to a *suppression* of conscious awareness^64,65^ - the opposite effect as reported here. It is generally thought that different types (types, tokens) of visual information lead to these differences between repetition blindness and priming^64,66^. One potential explanation for the difference in outcome from the present results, is that repetition blindness is generally found by presenting words which form a sentence and are presented at a relatively slow rate^64,65^. It is possible that using these very complex stimuli, which require sentence comprehension, uses different brain mechanisms than for the stimuli used in the current work. It would be interesting to repeat the style of current tasks and analysis while using more complex stimuli and determine where the effects reverse.

Repetition blindness has also been found to occur even when simple alphanumeric letters are used as targets^67,68^. This study found the AB was largest when the targets repeated in identity (i.e., the same letter) compared to when the targets were different letters. While these results superficially contradict the current findings, the subtle differences in methods may have been critical. In all the current experiments, the target identities never exactly repeated (orientations were at least 1° different or the same letter or number was not used for both targets). Instead, of classifying trials are repeated or not, as is convention in repetition blindness studies, we instead used the graded similarity between targets. It is possible that we may have seen suppression for targets that *exactly* repeated in identity, but still the enhanced for *similar, but not identical*, targets.

While the current work only focuses on the role of targets, properties of the distractors are also known to influence the ability to bring the post-T1 targets to conscious awareness^42^. One particularly powerful demonstration of this effect showed participants can report three sequential targets, whereas if the middle item is a distractor, they miss the final target^69^. Our previous work^20^ has shown that attention only boosts the neural representation of targets while distractors are unaffected, suggesting an early filtering of non-target features. Importantly for the current work, this sparing from filtering occurred regardless of the similarity between the two targets. We have previously shown that the perceived orientation of the target is also attracted towards the orientation of the following distractor, in addition to the other target^19^. These results collectively suggest that priming for the T1 information which is selective only for targets is established when there has been one distractor presented. Without the distractors, it is possible the system does not need to boost the gain of the T1 representation and therefore no target filtering occurs. It would be therefore interesting to test these similarity effects using a skeletal RSVP paradigm consisting of only two targets which reliably induce an AB^70^.

The increased in accuracy of similar targets reported in Experiment 1, along with the attractive biasing of target orientations are reminiscent of serial dependency effects^28,29^ and opposite to tilt-aftereffects where adaptation to an orientation causes repulsion for similar orientations and decreased accuracy^71,72^. The repulsive tilt aftereffect is well predicted by simple information channel models^72,73^, which are consistent with the neurophysiological evidence that visual neurons exhibit orientation-selective adaptation with the most similar orientations causing the greatest adaptation^74,75^. Using the same simple model architecture, but having the adapter increase the sensitivity of the affected channels allows for attractive aftereffects which are consistent with serial dependency^72^. This increase in sensitivity is consistent with the increase in orientation-selectivity when attention is directed towards stimuli^23,76^. These results can, therefore, be explained by a channel-based model where attention causes a boost in the representation to the orientation of the first target.

## Methods

### Participants

#### Experiment 1 and 2

As previously reported for this dataset^20^, for Experiment 1, 22 (13 females, 9 males) participants between the ages of 19 and 33 years (median age = 22 years) were recruited from a paid participant pool and reimbursed at AUD$20/hr. For Experiment 2, 23 (14 females, 9 males) participants between the ages of 19 and 33 years (median age = 23 years) were recruited from the same pool. Each person provided written informed consent prior to participation and had normal or corrected-to-normal vision. The study was approved by The University of Queensland Human Research Ethics Committee and was in accordance with the Declaration of Helsinki.

#### Experiment 3

As previously reported for this dataset^38^, 48 (26 female, 22 male) participants between the ages of 17 and 53 years (median age = 18 years) were recruited from the University of Western Australia psychology participant pool and given partial course credit. Each person provided written informed consent prior to participation and had normal or corrected-to-normal vision. The study was approved by the University of Western Australia Human Research Ethics committee and was in accordance with the Declaration of Helsinki.

#### Experiment 4

119 (68 females, 51 males) participants between the ages of 17 and 43 years (median age = 19 years) were recruited from the University of Western Australia psychology participant pool and given partial course credit. Each person provided written informed consent prior to participation and had normal or corrected-to-normal vision. The study was approved by the University of Western Australia Human Research Ethics committee and was in accordance with the Declaration of Helsinki.

#### Experiment 5

Three (two females) mice (C57BL/6rl) were used in this experiment. The mice were housed in a ventilated and air filtered climate-controlled environment with a reversed 12-hour light - dark cycle. The mice were between 7 and 9 weeks old at the time of recordings. The methods were approved by the Australian National University Animal Experimentation and Ethics Committee (A2019/11).

### Tasks

#### Experiments 1 and 2

The RSVP stream consisted of 20 Gabors (~ 5**°** diameter, 0.71**°** standard deviation), each presented for 40 ms with an 80 ms blank interval between stimuli, giving a presentation rate of 8.33 Hz. The participant was required to reproduce the orientation of the two higher-spatial frequency target Gabors (2 c/°) while ignoring the lower spatial-frequency distractors (1 c/°) in the order they were presented. Following the presentation of the final Gabor, a black circle appeared with a yellow line in the centre. The participants controlled the orientation of the line using the mouse and clicked when they were satisfied with the response. The participants were shown a feedback screen after making both responses for 600 ms before the next trial began. In both experiments, there were 600 trials equally distributed across the Lags (1, 2, 3, 5, 7 in Experiment 1; 3, 7 in Experiment 2). Between 4 and 8 distractors were presented before the first target. For Experiment 2, we recorded EEG using a BioSemi64 system while participants completed the task. The details for recording and pre-processing are reported in Tang, et al.^20^.

#### Experiment 3

The RSVP stream consisted of 10 items each displayed for 100 ms and followed immediately by the next item, giving a 10 Hz presentation rate. The targets consisted of the digits 2-9 and distractors were English upper-case letters, except B, I O and Q. The data was drawn from the assessment condition from the original paper^38^ which contained Lag 2 and 7 trials. There were 100 trials equally distributed across the two Lags. Participants initiated the RSVP by pressing the space bar and were prompted to type the identity of the targets after the presentation after which they could initiate the next trial.

#### Experiment 4

The RSVP stream consisted of 10 items each displayed for 30 ms with a 70 ms inter-stimulus interval, giving a presentation rate of 10 Hz. The targets were English letters, except I, O, Q, Z which were omitted due to their structural similarity with numbers^43^. The distractors were the digits 0 through 9 or pseudo-letters which are geometric shapes with a letter-like appearance^77^. The different types of distractors were randomly intermixed across trials. There were 4 to 8 distractors before T1. The identity of the targets and distractors was chosen pseudo-randomly with the previous that targets could not repeat in one trial. Each trial began with the presentation of a fixation cross with the participants initiating the RSVP by pressing the space bar. Participants were required to type the targets in the order they were presented. Following the response, the fixation cross re-appeared and the participants could initiate the next trial when they were ready. There were 400 trials in total, equally distributed across the lags (1, 2, 3, 5, 7).

#### Experiment 5

The mice initially had a surgery to implant a head bar, which allowed stable extracellular recordings, and for a craniotomy. For the surgery, the mice were initially anesthetized using 4% isoflurane delivered at 0.6 - 0.8 L\min. Following loss of consciousness, the animals were moved to a heated blanket (maintained at 37°, Physitemp Instruments) and were placed in a custom-designed stereotaxic head mount, with isoflurane delivered between 1 and 2% to maintain study anaesthesia. Consciousness was monitored throughout the surgery using foot pinches. The scalp and fascia over the dorsal surface of the skull were removed to allow for the headbar to be implanted and craniotomy to be made. Hydrogen peroxide (5%) was briefly applied to clean the site and to dry the bone. Headbars (International Brain Laboratory) were then attached to the scalp using UV-cured dental cement. Following this, a 3 mm circular craniotomy was made of left visual cortex (centred 3 mm lateral and 2 mm anterior to Lambda), which was then covered by a glass cover slip to protect the brain.

Following recovery, the mice were handled for one day and allowed to freely explore the recording setup for 30-60 mins for two days. Mice were then habituated for head fixation in a custom-designed head bar holder for 1 to 2 weeks. The mice were free to run on a circular disc that could freely rotate, with the movement captured by a rotary encoder. After the mice could tolerate head fixation for ~70-90 mins, the cover slip over the craniotomy was removed during a brief (< 20 min) surgery under isoflurane and with the first recording starting 4-5 hours later. Recordings were made 5 days a week for 1 week.

The stimuli were presented using PsychToolbox on a 9.7-inch monitor (LP097QX1) on the right of the animal, angled ~30º parallel to their body. The target stimuli were the same letters used in Experiment 4, consisting of all English letters except, I, O, Q, and Z. The letters presented in the centre of the monitor and increased in size to extend over ~50º of visual angle. During 20-item RSVP, a pseudo random sequence, drawn without replacement, of letters was shown. Each item was presented for 33 ms with a 66 ms interstimulus interval. A 1-1.1s inter-trial interval separated each of the 100 RSVP streams presented during the experiment. The mouse passively viewed the items and was not required to complete any task. Before the experiment, a high-density (960 channels, 384 active electrodes) Neuropixel probe^78^ was inserted using a micro-manipulator (www.newscale.com) into the left primary visual cortex. The activity digitized at 30 kHz using SpikeGLX (www.github.com/billkarsh/SpikeGLX) and synced to visual stimulus presentation using a photodiode attached to the computer monitor. Recordings were spike sorted using Kilosort 3^79^ with the results manually curated using Phy (https://github.com/cortex-lab/phy).

### Analysis

#### Experiment 1 and 2

Targets were marked as correct if the participant could reproduce the orientation within 30° of the presented orientation. In the initial analysis (Figure CF), similar trials were defined as those which the orientations of the two targets were within 30° of each other. For the later analyses (Figure D), accuracy was binned in from −90° to 90° in 20° steps of the difference between T1 and T2. To quantify the tuning, a Gaussian (Equation 1) was fit to the accuracy scores using non-linear least square:

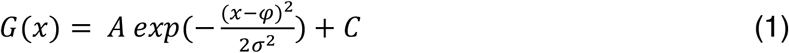

Where *A* is the amplitude, reflecting the number of responses around the reported orientation, *φ* is the orientation on which the function is centred (in degrees), *σ* is the standard deviation (degrees), which provides an index of the precision of participants’ responses, and *C* is a constant used to account for changes in the baseline rate. regression.

Following the original analysis^20^, the bias in accuracy was measured in a similar manner to accuracy (Figure E). For each trial, the mean orientation error (the difference between presented and reported orientation) was found for each bin in the difference between T1 and T2. To quantify the bias, we fit first derivative of Gaussian functions using non-linear least-squared regression:

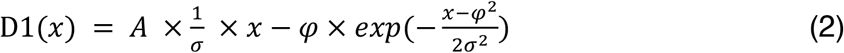

Where *A* is the amplitude, *φ* is the orientation on which the function is centred (in degrees), and *σ* is the standard deviation (degrees).

#### Computational model of temporal processing

The model of orientation processing was based on previous work examining orientation and motion-selective adaptation^72,73,80^. The model consisted of six Gaussian orientation-selective “neurons”, maximally sensitive to orientations from 0° to 150° in 30° steps, each with a standard deviation of 25° (Figure 1A). The probability distribution of the Gaussian reflects the neuron’s sensitivity to the presented stimuli. At the beginning of each trial, all the neurons are equally sensitive, and the population is therefore equally sensitive to all orientations. To model the effect of attention, the sensitivity of the population is modulated by 1 minus the response to T1 which is multiplied by a gain factor (set as 0.65) to scale the magnitude of changes. These modulated channels are used to then find the response to T2. To quantify the response to T2 is fit with the Gaussian function in Equation 1 with factors A and sigma being used to measure magnitude of response and decoded orientation, respectively. To determine the how orientation T1 biased the perceived orientation of T2 (Figure 1C), the presented orientation was subtracted from the decoded orientation. To determine how the orientation of T1 affected the magnitude of the response to T1, which is serving as a proxy for accuracy, the magnitude of the response was used.

#### Forward encoding modelling

To determine whether the computational model where T1 causes suppression of unattended representation, we used a forward (or inverted) encoding modelling approach to EEG activity. This approach determines patterns of EEG activity selective for feature-based representation, in this case orientation, information using regression. Typically, in the approach^20,23,31–33^, the regressors for any one orientation (e.g. 0°) are always the same regardless of the trial type. The regression matrix is then used to determine the patterns of activity selective for the presented orientations by regression against the measured pattern of neural activity. For the current work, we used a modified approach to forward encoding modelling to determine which model of orientation processing was the best fit for the recorded EEG activity. To do this, we used the computational model to produce the regressors for T2, which can be modified on a trial-by-trial basis by the orientation of T1. The gain parameter affected how T1 affects T2. A gain value of 0.65 was used for the priming model which decreases sensitivity of the orientations T1 did not activity. A gain value of −0.65 was used for the suppression model which enhances sensitivity of the orientations T1 did not activate. For the no priming models, a gain value of 1 was used which stops T1 affecting the response to T2.

The different regression matrixes for the three models were used to determine orientation selectivity within the pattern of EEG activity by solving the linear equation 3.

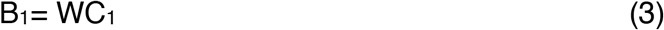

where B_1_ is the electrode data (64 channels × N training trials) for the training data, C_1_ (6 neurons/channels) is the regression matrix for the training data and W (64 electrodes x N test trials) is the weight matrix for the estimate sensors. Following previous work ^19,27,76^, we separately estimated the weights for each EEG electrode using least square regression to solving Equation 4:

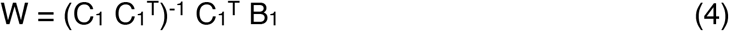

In line with previous work^20,33^, we removed the correlation between EEG electrodes which can inhibit finding the correct linear solution. To do this, we estimated and removed the noise correlation between electrodes using regularization^81^ by dividing W. The channel response in test set C_2_ (6 neurons/channels × N test trials) was estimated using the weights in (4) applied it to activity in B_2_ (64 electrodes × N test trials) as per Equation 5.

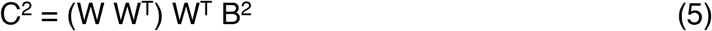

We used a 10-fold cross-validation approach to avoid overfitting, where X-1 trials were used to train the model which was tested on the remaining (X) trials. The process was repeated until all trials had served as both test and training sets. The whole procedure was repeated for every time point in the trials (−100 ms before stimulus to 500 ms after). So the trials could be aligned across different orientations, we reconstructed the item representation^20,23^ by multiplying the channel result with the unbiased orientation model (180 orientations × N time × N trials). We averaged the trials for each model and used a filtered the reconstructions over time using a 32 ms Gaussian kernel^20,32,33^. To quantify the orientation encoding over subjects, we followed previous work^20,32,33^ by fitting the smoothed reconstructions with a Gaussian function (Equation 1) with the amplitude of the Gaussian reflecting orientation selectivity.

### Mouse neuronal recordings in primary visual cortex

The spiking data were epoched relative (−0.5 to 1 s in 10 ms bins) to the presentation of the stimulus. Both single and multi-unit activity, regardless of responsiveness to the stimulus, were included in the analysis. Multivariate linear decoding was used to determine the population selectivity to the letters. To do this, linear discriminant analysis was to predict which letter was presented on each trial from the neuronal recordings. We used the same cross-fold validation procedure which was applied at each time point. As spiking data is relatively sparse, we smoothed the data with a temporal Gaussian (50 ms sigma) to allow better decoding. Decoding accuracy was determined by the predicted trial label matching the actual label. Decoding was conducted on the simultaneously-recorded population data for each session.

We used Representational Similarity Analysis^50,51^, to determine the representational similarity of the population response in V1 neurons across the stimuli. To do this, we used all units/neurons recorded on the top 1500 um of the recording shank to only use those in visual cortex. We found the average response across all presentations of each letter and took the average spike count from 100 to 300 ms after stimulus presentation which corresponded with the best decoding time from the multivariate analysis. We used 2510 units recorded across seven sessions to perform the RSA, which was given by correlating the average population response with each letter with all other letters. Following standard procedures, we used the dissimilarity (1 – r) as the metric. To determine whether similarly presented letters were more likely to be able to be consciously reported by the human participants, we found the difference in dissimilarity between the two targets.

#### Deep neural network analysis

We used a AlexNet^52^, a DNN, to determine the low-level similarity of the letters in a manner consistent with the visual system. We choose as AlexNet because it has previously been shown to be consistent with well-known aspects of the primate visual system^53,55^ and has a relatively simple architecture. The pre-trained network was implemented in MATLAB 2021a. AlexNet is organised in 8 major layers which feed hierarchically forward to the preceding layer (Figure 5A). The first five layers are convolutional layers (*conv1, conv2, conv3, con4, conv5*) which are organised in feature height x feature width x feature map channels. The final stages of the model are fully-connected layers (fc6, fc7) with a 1D structure of neurons which are all connected. The network has pre-trained on 1.2 million labelled natural images to have 1000 object categories. We presented the network with images of the letter stimuli used in the RSVP and found the activation for all the neurons within each layer. The same RSA approach was used as in the mouse V1 recordings.

#### Statistics

Any NaN value was replaced by the participant’s average for that comparison for all analyses in the work. Sphericity was tested for all ANOVAs, and if violated, a Greenhouse-Geisler correction was applied. Bonferroni corrections were used for all multiple comparisons. Fieldtrip toolbox^82^ was used to calculate the 2D cluster-permutation (n = 1,500) testing for the group level effects of EEG to determine whether the time course of topography difference between the conditions. For the time-resolved analysis (Figure 4D), cluster-based permutation testing was used to correct for multiple comparisons over the time series, with a cluster-form threshold of *p*<0.05 and significance threshold of *p*<0.05 used to determine whether there were differences between conditions.

## Data and Code availability

The data for Experiments 1 and 2 are available at: https://osf.io/f9g6h. The data for Experiments 3, 4 and 5 are available at: https://osf.io/unvyj. The code associated with the paper is available at: https://github.com/MatthewFTang/AttentionSelectivityTemporalDynamics.

## Acknowledgements

This work was supported by the Australian Research Council (ARC) with a Discovery Early Career Award (DE210100508) to MFT, the Centre of Excellence for Integrative Brain Function (CE140100007) grant to JBM and EA, an Australian Laureate Fellowship (FL110100103) to JBM, and Discovery Projects to TAWV (DP120102313) and EA (DP170100908). MFT, JBM and EA were also supported by an NHMRC project grant (APP1165337). JBM was also supported by the Canadian Institute for Advanced Research (CIFAR). JTE was supported by a Discovery Grant from the Natural Sciences and Engineering Research Council (Canada).

## Supplementary information

**Supplementary Figure 1.**
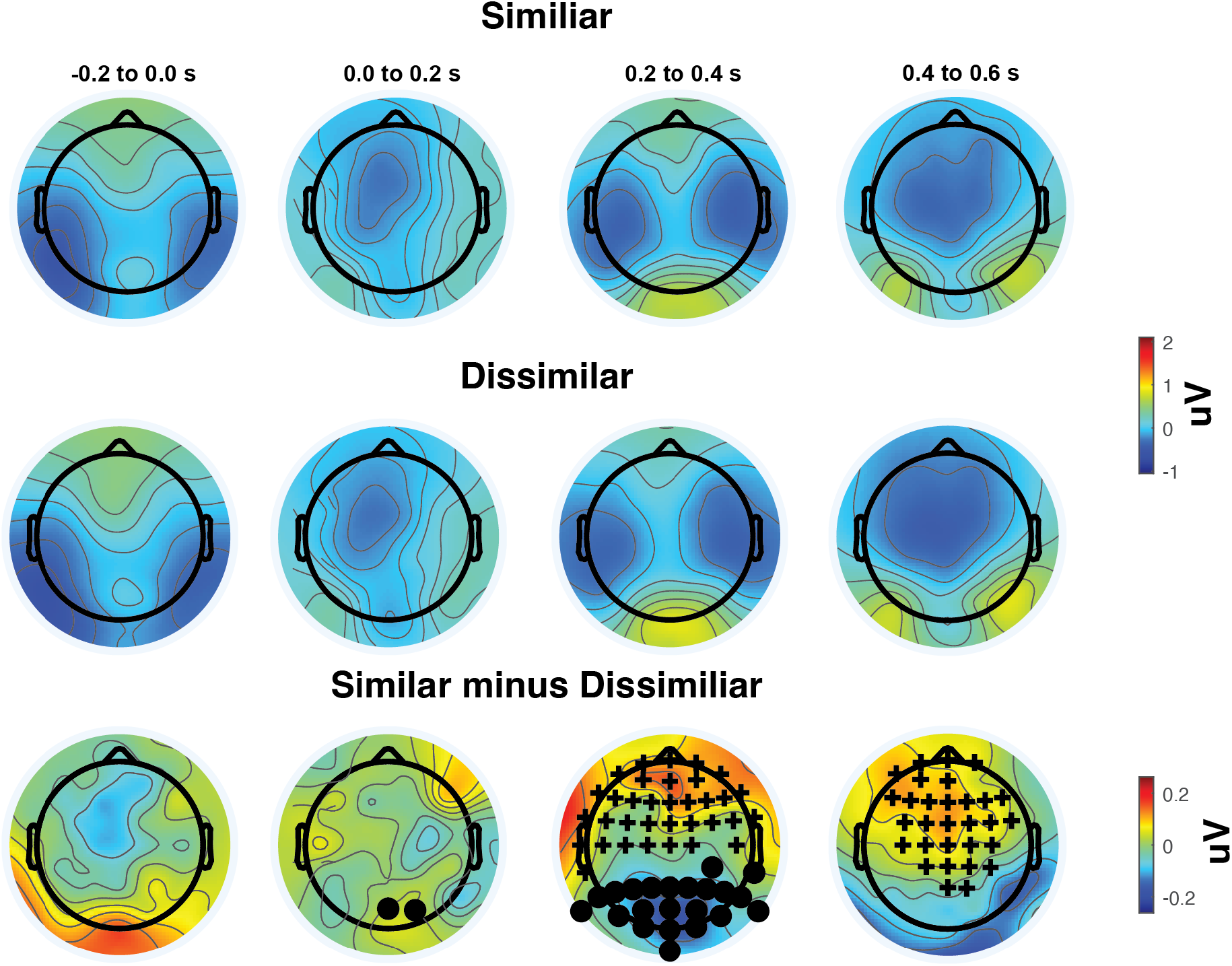
Topographic plots of EEG activity over time for Similar (top tow) and Dissimilar (middle row) stimuli in Experiment 2. The bottom row shows the difference between the conditions. The crosses indicate positivity cluster-permuted differences ^80^ between the conditions while the crosses indicate negative differences.

**Supplementary Figure 2.**
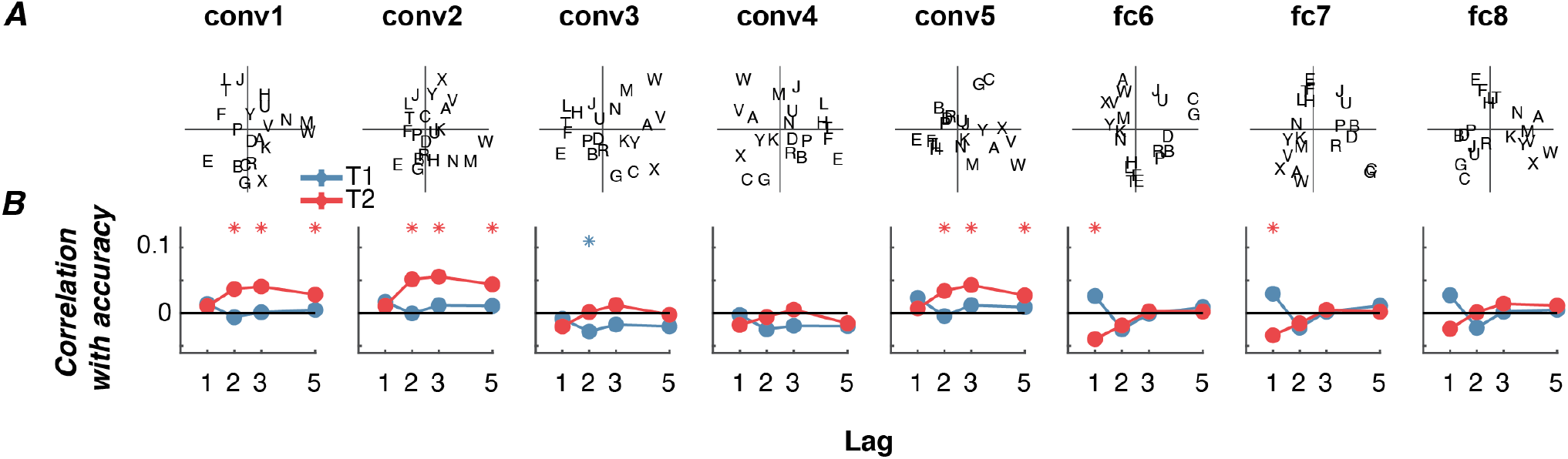
Using a deep neural network (AlexNet) trained on natural image classification to determine the effect of low-level similarity on target similarity effects in Experiment 4. (B) RSA visualised using multidimensional scaling for each layer. Similarly-represented letters are shown close together. Examining Conv-1 you can see in the top-left quadrant there’s letters while a single vertical line (I, T, F), the top-right quadrant has letters with oblique angles (X, V, A), bottom left has letters with curves (G, B, D). (B) For each subject, the difference in order or representational similarity between the targets for each of the different AlexNet layers was found for each trial and correlated with T1 and T2 accuracy to determine whether it predicts the detection for each different lag and target. Asterixis show Bonferroni-corrected one-sample t-test showing correlation is greater than 0, calculated separately for each target. The error bars indicate ±1 standard error.

